# Novel human monoclonal antibodies with enhanced sensitivity for lipoarabinomannan antigens present in urines of TB patients

**DOI:** 10.64898/2026.06.28.735056

**Authors:** Alok Choudhary, Deendayal Patel, William J. Honnen, Afsal Kolloli, Charles Reichman, Kamaljit Kaur, Ruixiang Blake Zheng, Elizabeth Nakabugo, Nasinghe Emmanuel, Lydia Nakiyingi, Todd Lowary, Abraham Pinter

## Abstract

Lipoarabinomannan (LAM) is a useful biomarker for detection of *M. tuberculosis* infection and disease. Related antigens can be detected in urine samples of TB patients by combinations of monoclonal antibodies (mAbs) directed against specific epitopes expressed in LAM. While sensitive for samples from patients with active TB disease who have HIV-1 co-infections, these assays are less effective for other populations, and there is therefore a need for more sensitive antibodies that can improve the sensitivity of these assays. Here we characterize the antigen and epitope specificities, sequence diversity and isotype dependencies of eight LAM-specific human mAbs that target five distinct arabinose- and mannose-dependent epitopes present in LAM and lipoarabinomannan (LM). Whereas all of the mAbs recognized ManLAM, only a few, including A194-01, consistently detected antigens in TB+ urine samples. Converting A194-01 from the IgG1 to the IgM isotype resulted in broader recognition of poly-Ara glycan epitopes, and increased sensitivity for clinical antigens when combined with several capture reagents, including RU95-C1, a novel antibody targeting the mannan domain of LAM. These results define novel epitopes that are differentially expressed in bacterial and urinary forms of LAM, and identify novel antibody combinations which possess enhanced diagnostic utility for clinical forms of LAM.

## Introduction

There is a large gap in the detection of new tuberculosis (TB) infections, with about 30% of new cases not being diagnosed early after infection ^1^. This highlights the need for a low-cost Point-of-Care (POC) assay that has sufficient sensitivity and requires minimal infrastructure or specialized skills, that can be used in low resources facilities and areas. The lipoarabonomannan (LAM) lipoglycan is a major mycobacterial surface antigen, and LAM metabolites secreted from bacteria and infected cells are important biomarkers for *M.tb* infection and activation. A commercial lateral flow assay that detects LAM in the urine of TB-infected subjects (Alere Determine TB LAM Ag) has proven to be a useful POC assay in diagnosing TB disease in a subset of HIV-1-coinfected patients ^2–4^, and early detection and subsequent treatment associated with the use of this test resulted in reduced mortality in this population ^4–6^. As a result, this test was strongly recommended by the WHO for use in the diagnosis of active TB in HIV-positive people who display signs and symptoms of TB, and in patients with advanced HIV disease and/or a CD4 cell count <200 ^7,8^.

The identification of high affinity monoclonal antibodies (mAbs) against LAM has allowed improvements in the sensitivity and specificity of these assays ^9–11^, and have led to a more sensitive POC test, the Fujifilm SILVAMP TB LAM (FujiLAM) assay. This assay utilizes uLAM capture mAb S4-20, that recognizes an methyl-thio-Xyl*f* (MTX)-dependent S4-20 epitope present in the capping domain of LAM from *M.tb* and other slow-growing mycobacteria ^24,25,12^, and the detector mAb A194-01, specific for a broadly conserved Ara4/Ara6-dependent epitope. This test also incorporates a proprietary silver amplification step that further increases assay sensitivity ^13–16^. Despite its greater sensitivity, the ability of the assay to detect LAM in samples from patients without HIV-1 coinfections remained too low to meet the WHO’s requirements to serve as a global screening assay ^15,17^, and recent efforts have not identified new antibodies that significantly improve the efficacy of these assays ^18,19^.

The majority of available LAM-specific mAbs were derived by immunization of animals with LAM antigens purified from *M.tb* or *M.Leprae* ^20,21^ or from phage antibody libraries ^22–24^, with only a limited number of antibodies derived from humans infected with *M.tb* ^9,25,26^ (reviewed in ^27^). In this paper we describe a total of eight human mAbs induced by TB infection, including six novel antibodies, and describe their antigen and epitope specificities, examine the contributions of isotype and variable region sequence to binding activities, and explore their potential in immunodiagnostic assays for clinical samples of LAM.

## RESULTS

### 1. Antigen specificity of mAbs

The antigen specificity of the panel of human mAbs was initially determined by direct binding titrations against LAM purified from three mycobacterial species (*M.tb, M.smeg and M.leprae*) and LM from *M.tb* (Fig. 1a). Four of the mAbs (A194-01, RU83-A8, RU49-01 and RU49-02) were reactive with all three LAM species, but not with M.tb LM (Fig. 1a), indicating that their targets were located solely in the arabinan domain. Three of the mAbs recognized both LAM and LM; RU95-C1 recognized all four antigens, with lower affinity for LepLAM and *M.tb* LM, while RU28-01 and RU61-H5 possessed highest affinity for ManLAM, lower reactivity for LepLAM and *M.tb* LM, but did not recognize PILAM. P30B9 was highly specific for ManLAM, with only slight crossreactivity for LepLAM, and did not recognize either PILAM or *M.tb* LM.

**Fig. 1.**
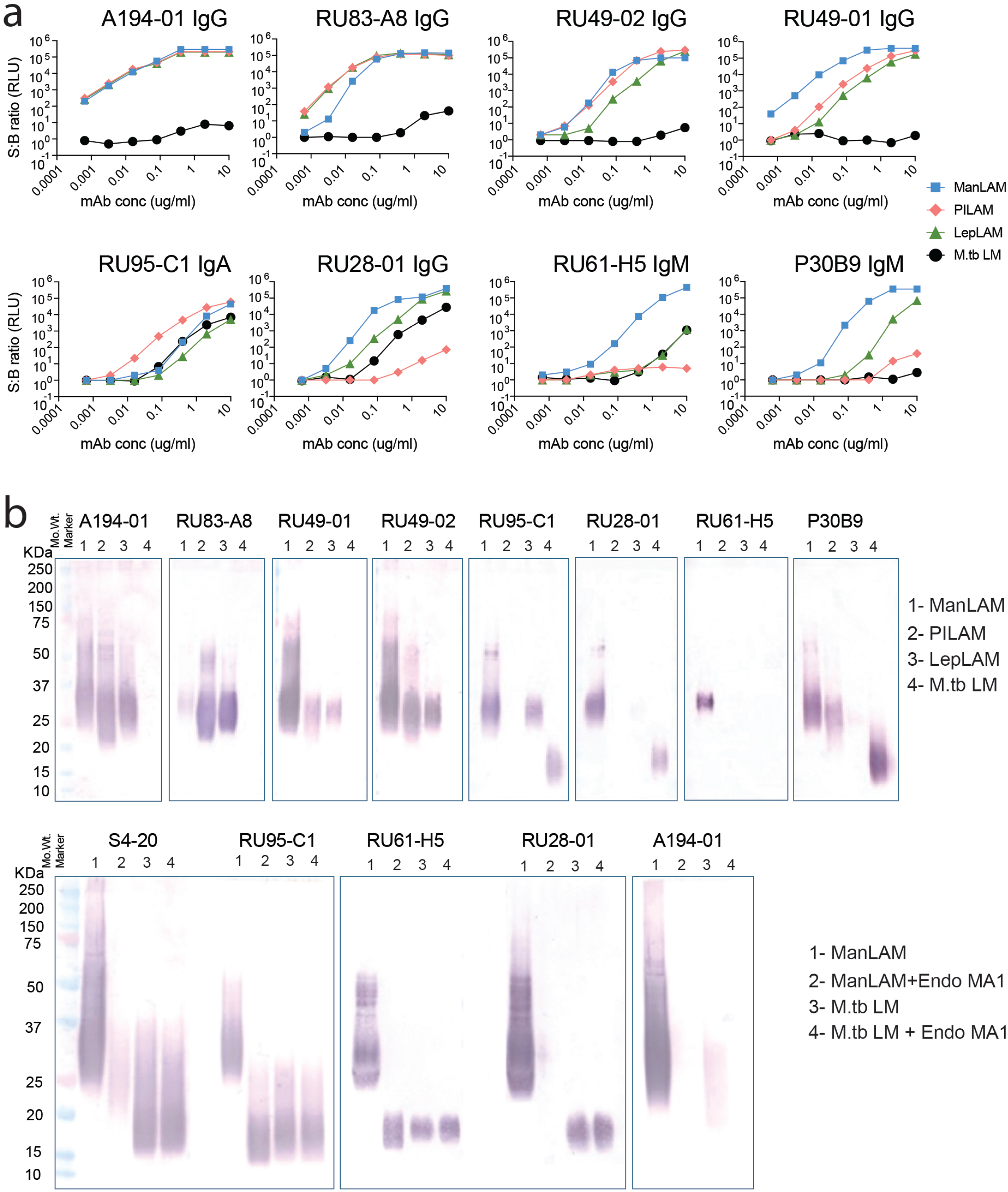
**a.** Antigenic analysis of eight human mAbs isolated from subjects with active TB infections. MAbs were titrated against purified mycobacterial lipoglycan antigens (LAM from *M.tb, M.Lep, M.smeg* and LM from *M.tb*) by ELISA, using a sensitive ECL-based readout. Fig. 1b- Confirmation of antibody specificities by Western blotting against the four antigens. The antibodies were stained with an HRP-labelled rabbit anti-human light chain secondary antibody. Fig. 1c- Comparison of reactivity of Man-dependent mAbs against *M.tb* LAM, LM, and the mannan domain generated by digestion of ManLAM with endoarabinase EndoMA1. The faint band seen in lane 3 for A194-01 running at the LAM position was absent after EndoMA1 digestion, and represented a minor contamination of the LM preparation by LAM.

Further analyses of these antibody specificities by Western blotting assays confirmed the relative preferences of the mAbs for the different LAM species, and the binding of RU95-C1, RU61-H5 and RU28-01 to the LM antigen (Fig. 1b). Differences in sensitivity for the three LAM antigens was more pronounced for several of the mAbs in the blots, presumably related to the lower sensitivities of the blots than the ELISA format. For example, the preferential recognition of RU83-A8 for PILAM over ManLAM was more evident in the blot than in the ELISA, and RU94-01 and RU94-02 showed a greater specificity for ManLAM over the other LAM antigens in the blots than in the ELISAs.

The basis for the recognition of LM by the three mAbs was further probed by comparing their reactivity in Western blots with LM, LAM, and the mannan domain generated by digestion of Man-LAM with Endo-MA1, an endo-α1-D-arabinofuranosidase cloned from *M.arabinogalactanolyticum* that digests α-(1→5)-linked arabinofuranosyl linkages and efficiently degrades the polyarabinose backbone in LAM ^28^. The MTX-Man_2_-dependent S4-20 and Ara-specific A194-01 mAbs were included as controls (Fig. 1c).

As expected, A194-01 recognized intact ManLAM, but not LM or the mannan fragment of endo-arabinase-digested LAM, and as previously reported ^12^, S4-20 reacted with both ManLAM and LM, but not with the mannan domain of LAM. RU28-01 resembled S4-20 in that it recognized LM but not the isolated mannan domain of ManLAM. In contrast, RU95-C1 and RU61-H5 both recognized all three antigens, indicating that their epitopes were present in the mannan domains of both LM and LAM. These different immunoreactivity patterns demonstrated the immunological distinctiveness and diversity of the mannan domains of LM and LAM.

### 2a. Epitope specificities of mAbs

To better understand the basis of these antigen reactivities, the epitope specificities of these mAbs were mapped by titrating their binding activities against a microarray containing 61 synthetic glycan structures that represented a large variety of carbohydrate motifs present in ManLAM and other mycobacterial lipoglycans ^9,29,30^. The glycans were conjugated to a BSA carrier, to mimic the multivalent presentation of these epitopes in the native LAM antigen. Structures of 32 key polyarabinose, arabinomannose and polymannose structures recognized by these mAbs are shown in Suppl. Fig. 1a, and the reactivities of the mAbs against these structures are shown in Suppl. Fig. 1b, and are described in greater detail below and summarized in Table 1. The glycoconjugate array analysis indicated that mAbs A194-01, RU83-A8, RU49-01 and RU49-02 recognized mannose-free polyarabinose structures, while RU61-H5, RU28-01, P30B9 and RU95-C1 were dependent on the presence of mannose-containing structures.

**Table 1.** Summary of antibody characteristics.

| Antibody | Isotype dependence | Ag Specificity | Epitope characteristics | VH chain aa mutation level | VL chain aa mutation level |
| --- | --- | --- | --- | --- | --- |
| A194-01 | None | ManLAM, PILAM, LepLAM neg. for LM | Strongest for uncapped Ara4/Ara6, weaker reactivities for Man1 caps, weak or no reactivity for larger Man caps | 10.6% | 12.5% |
| RU83-A8 | None | ManLAM, PILAM, LepLAM, neg. for LM | Uncapped Ara4/Ara6, no reactivity for structures with Man caps | 9.5% | 5.7% |
| RU49-01 | None | ManLAM > PILAM, LepLAM. neg. for LM | Poly-Ara- preference for truncated terminii with α-(1→3)-linked Ara branch | 10.3% | 7.9% |
| RU49-02 | None | ManLAM = PILAM > LepLAM, neg. for LM | Broad, Long chain poly-Ara | 15.8% | 10.2% |
| RU61-H5 | IgA/IgM | ManLAM >>LepLAM, neg. for PILAM, crossreactive with LM | Preferential reactivity for Man2 caps, weaker for Man1, and Man3 caps, crossreactive with LM | 6.3% | 2.3% |
| P30B9 | IgA/IgM | ManLAM, neg. for PILAM and LM, slight reactivity with LepLAM | Specific for Man2 caps | 16.0% | 5.7% |
| RU28-01 | None | ManLAM >>LepLAM, neg. for PILAM, crossreactive with LM | Preferential reactivity with Man3 caps, weaker reactivity, with Man1 and Man2 caps | 20.0% | 3.2% |
| RU95-C1 | None | PILAM > MAnLAM >> LepLAM, specific for LM and LAM mannan domain | α-(1→6)-linked Man trimer in mannan domain of LAM, and LM, inhibited by α-(1→2)-linked Man branching | 18.9% | 13.7% |

### 2b. Fine specificities of four mannose-dependent antibodies

ManLAM has two structural regions that contain mannose sugars, the capping region located at non-reducing ends of the arabinan domain, and the mannan domain joining the arabinan domain to the phosphoinisitol-lipid anchor. Mannose caps consisting of α-(1→2) -linked mono-, di-, or trimannosides are attached at the non-reducing end of the arabinan domain, with a limited number of the terminii further modified by attachment of α-(1→4)-linked 5’ MTX ^21,31–35^. The mannan domain consists of α1→6-linked mannose chains with short α1→2-linked mannose branches ^36,37^. Key structures present in both of these domains were used to define the fine specificities of the four mannose-dependent antibodies, with A194-01 included for comparison (Fig. 2a-b).

**Fig. 2.**
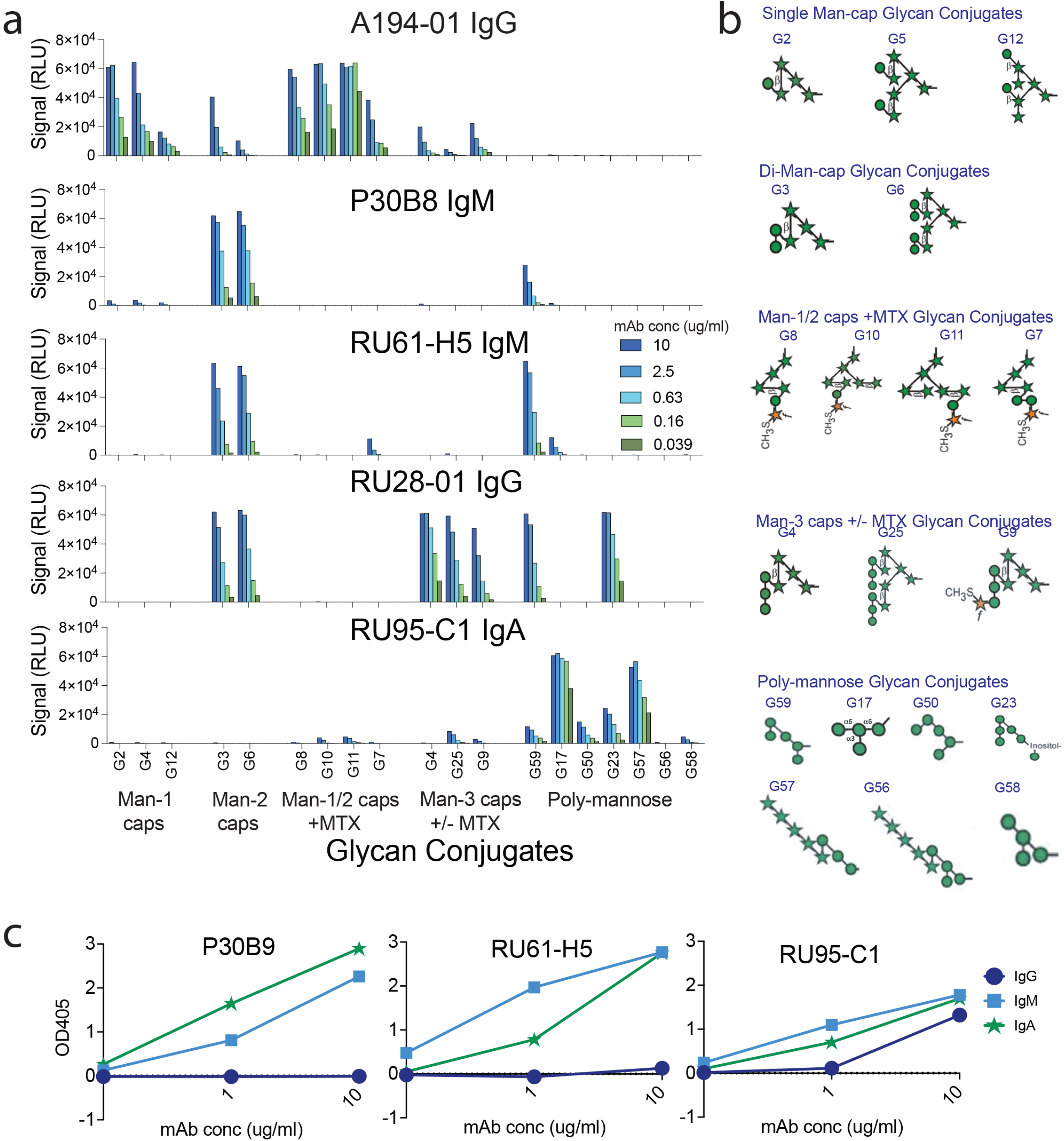
**a.** Binding titrations for A194-01 and four mannose-dependent mAbs against selected glycoconjugates. **Fig. 2b.** Structures of the mannose-containing glycans used in this experiment. **Fig. 2c.** Effect of isotype on function of mAbs naturally expressed in IgM form. The IgM constant domain of these antibodies was exchanged for that of either IgA or IgG, and the binding activities of the purified antibodies compared to that of the natural IgM isoforms.

P30B9 and RU61-H5 both reacted strongly with two dimannose caps linked Ara4 (G3) and Ara6 (G6) polyarabinose structures, and binding of both antibodies was reduced by the attachment of MTX (G7), indicating that an unmodified terminal mannose sugar was a preferred interaction sit-binosr structures. RU61-H5, and to a lesser extent, P30B9, also recognized G59, a pentamannose glycan consisting of a α-Man-(1→6)-α-Man-(1→6)-α-Man trisaccharide with α-Man-(1→2) branches at the two ends. These dimannose branches resembled the Man2 cap structure, presumably accounting for the cross-reactivity between the caps and the poly-mannose structures. The cross-reactivity of these mAbs for the dimannose glycan present either in the cap or polymannose structures suggested that the dimannose sugars were the dominant target, and that the arabinose backbone was not essential for recognition, but instead may provide a framework for multivalent presention of these epitopes that facilitates recognition. Related dimannose motifs are also present in the mannan domains of both LAM and LM, and the recognition of these structures by RU61-H5 is consistent with the stronger crossreactivity of this antibody to this poly-mannose antigen.

RU28-01 did not recognize any of the uncapped or Man1-capped structures, and reacted with strongest affinity with structures containing α-(1→2)-linked trimannose structures presented on either polyarabinose (G4) or polymannose (G23) scaffolds. G23 is an inositol-containing hex-amannose structure that modeled the PIM6 glycan. RU28-01 bound with weaker affinity to Man2-capped Ara4 and Ara6 structures (G3 and G6), and to the pentamannose G59, which contained two terminal α-Man-(1→2)-α -Man branches that resembled the Man2 cap structure. Unlike the Man2-dependent mAbs, RU28-01 retained considerable affinity for an MTX-substituted Ara4Man3 (G9), but did not bind to the MTX-substituted Ara4Man2 (G7), indicating that while the MTX substitution abrogated the weaker Man2 interaction, it only partially inhibited the stronger interaction with the Man3 structure.

RU95-C1 was unique in that it recognized a number of the poly-mannose structures, but did not bind to any of the Ara-linked mannose cap structures. The common feature of the reactive glycoconjugates was the presence of an α-Man-(1→6)-α-Man-(1→6)-α-Man trisaccharide, characteristic of the core of the mannan domain. The strongest reactivity was with G17, which contained this trisaccharide with an α-(1→3)-linked Man branch off of the middle Man residue, and with G57, which contained the same trimannose core modified with an α-(1→5) linked Ara5 chain via α-(1→2)-linkage to the terminal mannose. Related structures (G59, G23, G50) containing an α-(1→2)-linked or an α-(1→3)-linked mannose branch off the terminal sugars of the α-Man-(1→6)-α-Man-(1→6)-α-Man trisaccharide had lower reactivities, while the presence of an α-(1→2)-linked Man branch on the middle Man residue of the trisaccharide (G56, G58) prevented recognition by RU95-C1. This data suggests that the minimal RU95-C1 epitope consisted of an α-(1→6)-linked trimannose structure, and that binding affinity was modulated in various ways by the presence of α-(1→2) and α-(1→3)-linked mannose branches.

### 2c. Role of antibody isotype in antibody function

Three of the eight mAbs (P30B9, RU61-H5, and RU95-C1) were isolated and expressed as IgMs, while the other mAbs were cloned in their native IgG isoform. The importance of isotype towards the binding activities of the native IgM mAbs was explored by exchanging the Fc domains of the IgMs for IgG or IgA domains. Conversion of P30B9 and RU61H5 IgM to the IgG isotype resulted in loss of all binding activity, while conversion to the IgA isotype resulted in increased binding activity (Fig. 2c). This indicated the affinities of the dimannose-dependent bivalent mAbs towards their targets in ManLAM are low, and the higher valency of the IgA and IgM forms was required to attain sufficient avidity to recognize these targets. For RU95-C1 all three isotypes remained active, although the IgG form possessed lower reactivity that either the IgA or IgM forms. For this reason, further experiments with RU95-C1 were performed with the IgA form, which was easier to express and purify than the IgM form.

### 3. Sequence analyses of anti-LAM mAbs

The genetic sequences of these mAbs were analyzed in order to obtain further insight into their nature and origin. A summary of their gene usages, level of somatic mutations and genetic homology is shown in Suppl. Table 1, homology trees for the H chains and L chains are shown in Suppl. Figs. 2a and b, respectively, and alignments of the VH and VL gene sequences with the nearest germline sequences are shown in Suppl. Fig. 3. With the exception of A194-01 and RU83-A3, the mAbs were derived from diverse VH and VK gene usage. The VH3 gene family was the most common, used by five of the eight antibodies, followed by two VH4 genes and a single VH1 gene. Seven of the eight light chains were derived from kappa genes, while one (RU49-01 mAb) used a VL1 lambda chain. VK3 genes were each used by three antibodies, VK1 by two antibodies, and VK2 and VK4 gene were each used by one antibody. The A194-01 and RU83-A8 sequences were closely related, and were derived from the same IGHV3-20*04 and IGHJ6*03 heavy chain, and IGKV-15*01 and IGKJ2*-01 light chain gene segments.

The presence of somatic hypermutation in both heavy and light chains suggests that these antibodies underwent antigen-driven affinity maturation. The mutation levels of the eight antibodies from their nearest germ-line sequences are shown in Suppl. Fig. 3 and summarized in Suppl. Table 1. These levels varied considerably for the mAbs, ranging from 6.3% to 20% for the IGVHs, and from 2.3% to 13.7% for the IGVL sequences. Variability in the VH-CDR regions ranged from 0 to 16.7% for the CDR1 domain, and 12.5 to 53.8% for the CDR2 region, with the highest levels of mutation due to the presence of five and six amino acid inserts in the CDR2 region of RU28-01 and P30B9, respectively. Mutation levels in the framework regions ranged from 0 - 11.8%.

### 4. Role of hypermutations and HCDR2 insertions on function of P30B9 and RU28-01

Contributing to the high mutation rates of the RU28-01 and P30B9 VH proteins (20.0% and 16.0% respectively) were their heavily mutated CDR2 domains, which included atypical insertions of five and six amino acids, accompanied by additional surrounding mutations (see Supp Fig. 2a, Suppl. Fig. 3). The importance of these insertions and mutations towards the function of these mAbs was explored by analysis of the effect of reversion to the germline sequences.

In addition to the six amino acid (aa) insertion in H-CDR2, P30B9 possessed nine substitution mutations in the FR2, CDR2 and FR3 domains. (Fig. 3a). Reverting the VH sequence back to the germ-line sequence while retaining the CDR3 region resulted in the complete loss of binding reactivity to ManLAM (Fig. 3b). Analyses of partial reversions revealed that the six aa insert was the more dominant requirement for function, because inserting the six aas into the GL revertant (GL + 6aa) resulted in complete recovery of reactivity, while deleting the insert from the wt sequence (WT – 6aa) resulted in a reduction in reactivity. As noted above, conversion from the IgM to the IgG isoform resulted in complete loss of ManLAM binding.

**Fig. 3.**
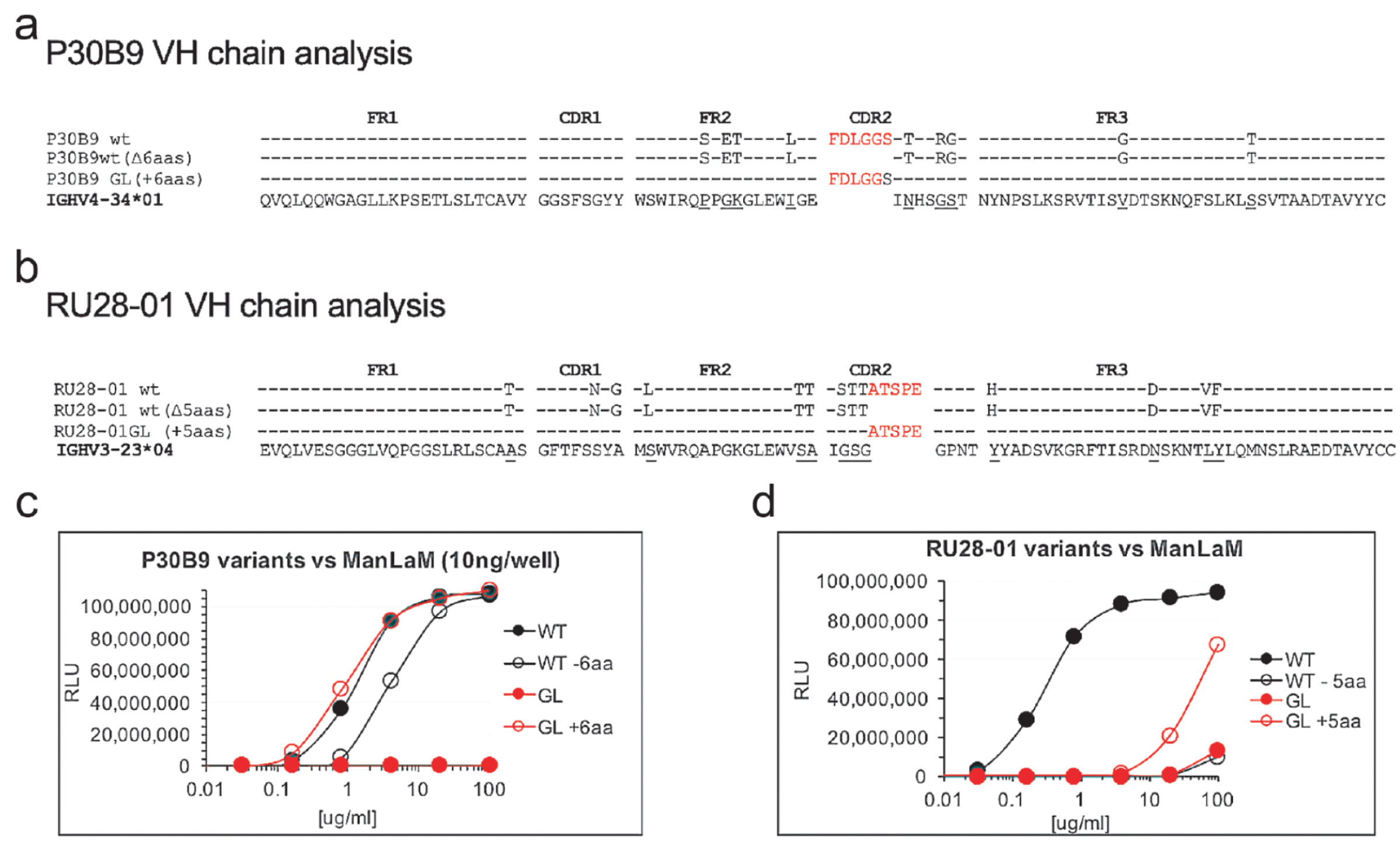
Homology of amino acid sequences of RU28-01 (Fig. 3a) and P30B9 (Fig. 3b) wt and mutant heavy chains with the nearest germline sequences. Binding titrations for the wt and variant forms of RU28-01 (Fig. 3c) and P30B9 (Fig. 3d) to ManLAM are shown.

RU28-01 possessed a five aa insertion in the VH-CDR2 region, and thirteen substitution mutations localized to the FR1, CDR1-FR2, and FR3 domains (Fig. 3a). Separately reverting these features to their germline sequences revealed that both the five aa insert and the somatic mutations were required for full function (Fig. 3c). While the GL sequence and the wt RU28-01 H chain mutant in which the five aa insert was deleted (WT-5aa) had identical low activities, insertion of the five aa insert into the GL seqence (GL+5aa) resulted in a partial restortion of function. This showed that for the RU28-01 mAb, the CDR2 insert was more critical than the somatic mutations for function.

### 5. Functional relationship between A194-01 and RU83A8

The genetic homology between A194-01 and RU83-A8 (Suppl. Table 1, Suppl. Fig. 3) suggested that these antibodies were members of a conserved family, which may possess related activities. A194-01 possessed similar affinities for ManLAM and PILAM, while RU83-A8 bound PILAM with similar affinity as A194-01, but had lower affinity for ManLAM (Fig. 4a). The functional homology between these mAbs was examined by generating chimeric antibodies in which the heavy and light chains of the two antibodies were swapped. Both the A194VH/83VK and 83VH/A194VK chimeras were active, and while they retained similar affinities to PILAM as the parental antibodies (Fig. 4b, top panel), the both had reduced affinities for ManLAM, which were intermediate between those of the parental antibodies (Fig. 4b, bottom panel). This indicates that the heavy and light chains of both antibodies are functionally interchangeable, and both contribute to their activity and specificity.

**Fig. 4.**
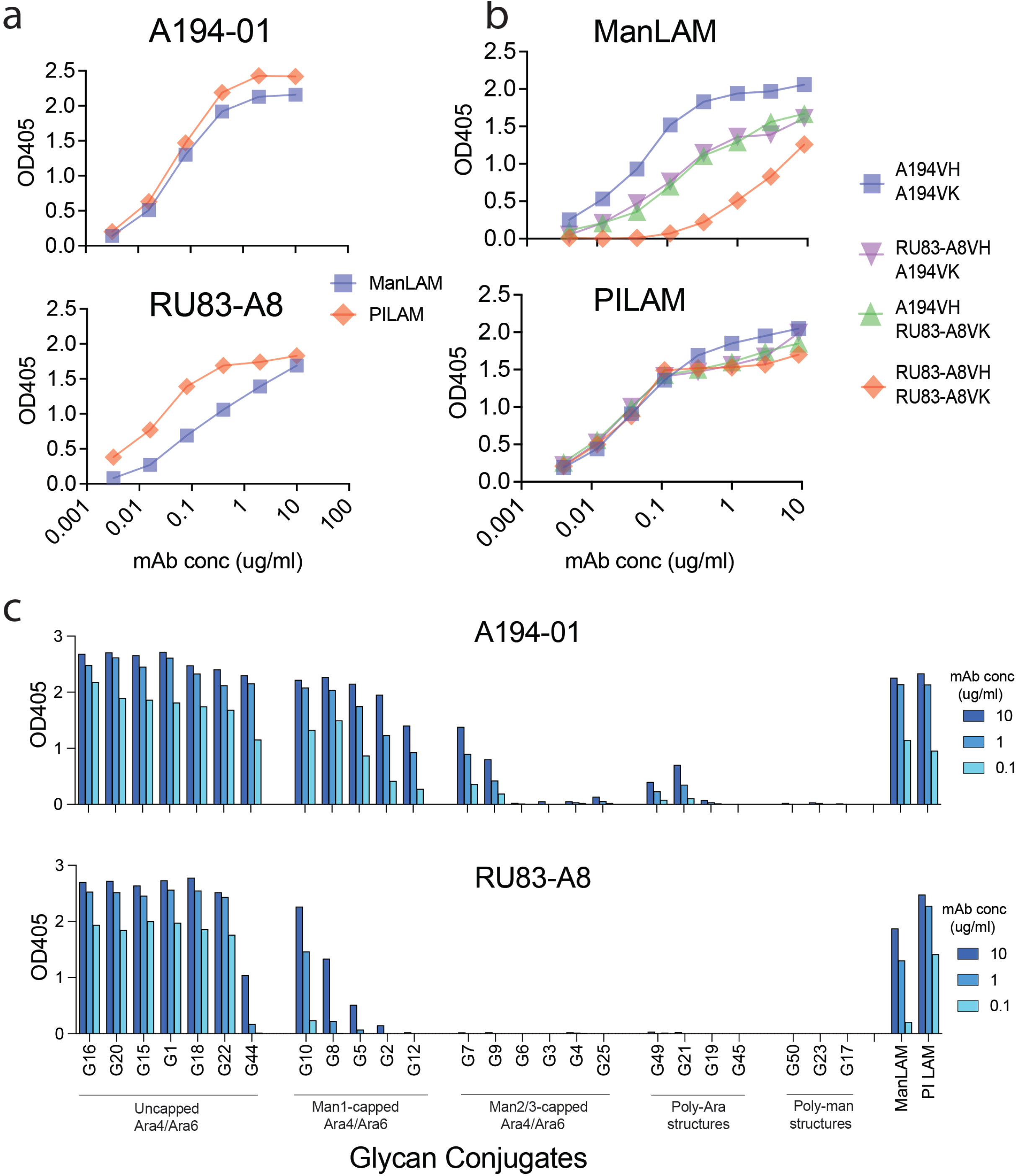
**a.** Binding titrations of A194-01 (top panel) and RU83A8 (bottom panel) were determined by ELISA for a selected set of synthetic glycoconjugates. **Fig. 4b.** Comparison of binding activities of A194-01 (top panel) and RU83-A8 (bottom panel) for ManLAM and PILAM. **Fig. 4c.** Binding activities of the two parental antibodies and their H and L chain chimeras for PILAM (top) and ManLAM (bottom).

The differences in affinity of the two mAbs for ManLAM and PILAM was explained by comparison of the fine epitope specificity of A194-01 and RU83-A8 (Fig. 4c, Suppl. Fig. 1b). Titrations of the binding activies of two mAbs against a selected subset of glycoconjugates shows that two mAbs possessed similar binding activities for a number of uncapped Ara-containing glycans containing either Ara4 and Ara6 motifs, and little if any reactivity with a series of larger poly-arabinose structures containing incomplete Ara4 or Ara6 terminii, indicating that the terminal Ara structures are critical targets for both mAbs. A194-01 possessed slightly reduced reactivity for structures containing Man1 caps, and bound more weakly to glycans containing larger Man2 and Man3 caps. This reduced affinity for Man-capped structures was magnified for RU83-A8, which weakly recognized structures capped with a single MTX-mannose motif, but did not bind to structures with the larger caps. The similar affinities to the uncapped Ara motifs explains the similar affinities to PILAM, and the reduced sensitivity of RU83-A8 to the mannose caps accounts for its reduced affinity for ManLAM.

### 6. Effect of A194-01 isotype on binding properties

The loss of reactivity of IgG isotypes of the natural IgM mAbs, P30B9 and RU61-H5, highlights the effect of higher valencies on affinity of antibodies that recognize the mannose caps of LAM. This raised the question of whether a higher valency might enhance the activity of a broadly reactive bivalent antibody such as A194-01. To test the effect of valency, the A194-01 VH and VL genes were cloned into vectors expressing different forms, ranging from monovalent Fabs and single-chain Fv (scFv), to bivalent and tetravalent constructs, and to decavalent IgMs. Initially, the ability of these different A194-01 forms to compete with the direct binding of the biotinylated A194-01 IgG for ManLAM was examined (Fig. 5a). Relative affinities were determined by comparing the molar concentrations of competing reagent required for 50% reduction of binding of the dimeric A194-01. The monomeric constructs possessed the lowest relative affinities (303 nM for theFab and 82 nM for the scFv), compared to 3.1 nM for the scFv dimer and 2.8 nM for the IgG itself. The tetrameric IgG-scFv competed more efficiently than the IgG (1.3 nM), while the pentameric IgM competed most efficiently (0.6 nM).

**Fig. 5.**
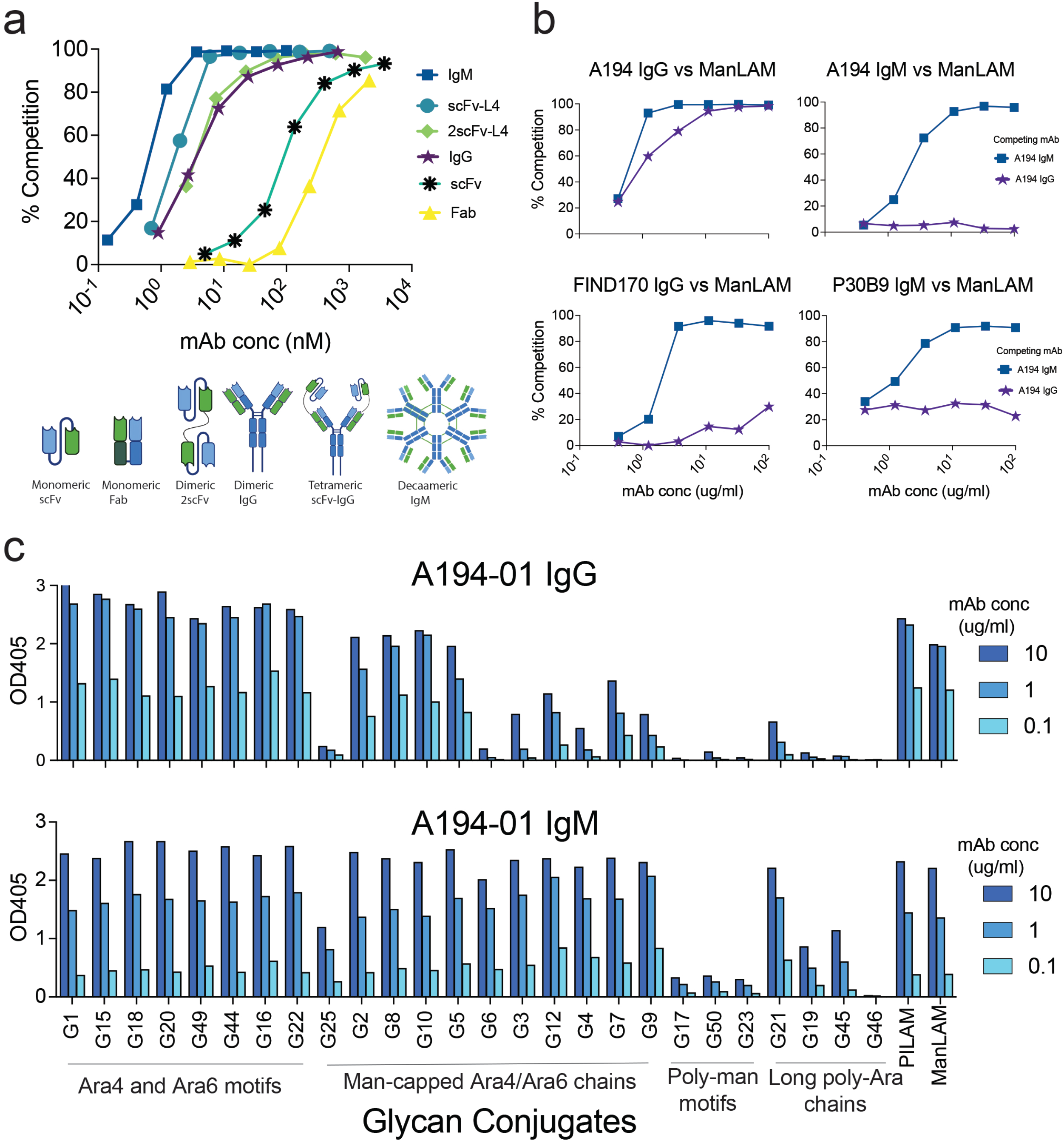
Effect of antibody valency on A194-01 affinity. a- Binding competition by direct ELISA of biotinylated A194-01 IgG by A194-01 isoforms with different valencies. b- Comparison of efficiencies of competition of A194-01 IgG vs. IgM forms for binding to ManLAM by anti-LAM antibodies with different epitope specificities. c- ELISA showing comparison of epitope specificities of A194-01 IgG and IgM forms against a subset of glycoconjugates described in Supp. Fig. 1 demonstrating the broader specificity of the IgM form.

The relative ability of the bivalent A194-1 IgG and pentavalent IgM to compete for binding of different antibodies to ManLAM was then measured (Fig. 5b). Both forms competed for binding of the A194-01 IgG isotype, with the A194-01 IgM competing more efficiently than the IgG form (upper left panel). However, only the IgM, but not the IgG, competed when the A194-01 IgM was used as the ligand (upper right panel). In addition, the IgM, but not the IgG form, competed for binding of two mAbs that recognized different LAM epitopes, FIND170 IgG, specific for capped and uncapped Ara6 structures, and P30B9 IgM, specific for Man2-capped structures (lower left and right panels). The limited ability of A194-01 IgG to compete with these two mAbs is consistent with data previously reported ^9^, and presumably reflects the efficient binding of FIND170 and P30B9 to mannose-capped structures which are abundant in ManLAM, for which A194-01 IgG possessed low affinity. However, the ability of the IgM form to efficiently compete for binding to these structures was unexpected.

To explain this potent competition, we compared the reactivity of the A194-01 IgG and IgM forms for a subset of the glycoconjugates used for mapping of the glycan epitopes recognized by the LAM-specific antibodies (Fig. 5c, Suppl. Fig. 1a). This assay showed that the A194-01 IgM recognized a broader range of glycan structures than the IgG form. Whereas the IgG form possessed low affinity for many of the mannose-capped structures, the IgM isotype consistently recognized all of these forms, and even reacted with several long-chain poly-Ara structures that did not possess the standard Ara4/Ara6 terminii. This increased binding breadth was consistent with the ability of the IgM isotype to compete for binding to antigens for which the IgG possessed low affinity.

### 7. The A194-01 IgM isoform possesses enhanced detection sensitivity for urinary LAM (uLAM)

The broader breadth and increased binding affinity of A194-01 IgM for the LAM-related glycans suggested that this isotype may also have advantages for the immunodetection of clinical forms of LAM. This was tested by comparing the detection sensitivities of A194-01 IgG vs IgM in capture assays for ManLAM and representative urine samples from six HIV-1 and TB-positive Ugandan subjects (Fig. 6a). A screening assay of all possible combinations of S4-20 and the eight mAbs described in this study revealed that while most of the antibody combinations allowed detection of ManLAM, only a limited number of combinations were able to detect the majority of the positive urine samples (Suppl. Fig. 4). We therefore compared the sensitivities of two of the broadly-reactive capture mAbs, S4-20 and RU95-C1, in combination with either A194-01 IgG and IgM, for the positive urine panel.

**Fig. 6.**
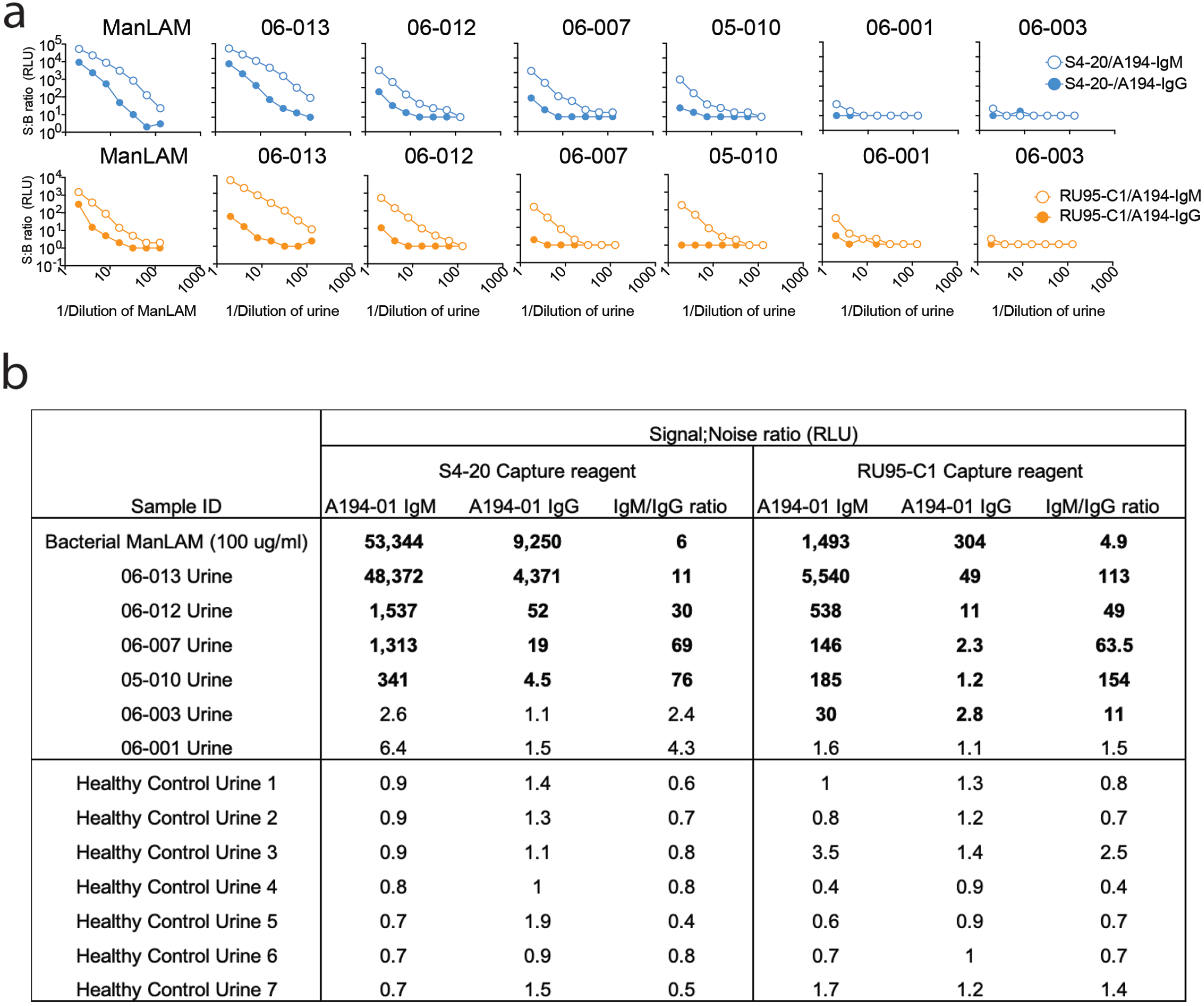
**a.** Enhanced sensitivity of A194-01 IgM for urinary LAM samples from subjects with microbiologically confirmed *M.tb* infections. Shown are capture assays performed with S4-20 or RU95-C1 capture antibodies and IgG and IgM forms of A194-01, against titrated urine samples from six Ugandan subjects co-infected with TB and HIV-1. **Fig. 6b**- Signal ratios of S4-20 and RU95-C1 capture and A194-01 IgM vs. IgG detection, for 1:2 dilutions of six positive and seven negative control urines.

For ManLAM and five of the six urines tested, the A194-01 IgM isotype consistently gave stronger signals and higher titers than the IgG isotype, and several urine samples that gave negligible signals with A194-01 IgG gave strong signals with the IgM (Fig. 6a). Although the S4-20 capture reagent was more sensitive than RU95-C1 when coupled with A194-01 IgG, the enhancement seen for A194-01 IgM was higher for RU95-C1 for several urine samples (Fig. 6b). The levels of enhancement with the IgM reagent for 1:2 diluted urines ranged from 7.8 to 53.5, with higher ratios seen for the samples that gave weaker signals with A194-01 IgG (Fig. 6b). When tested against a series of healthy control urines, both mAb combinations gave very low background signals, similar to that seen for the blocking reagent (2% BLOTTO) itself (Fig. 6b), indicating that the A194-01 IgM isotype increases sensitivity of detection of clinical forms of LAM, while retaining similar specificity as the IgG form.

### 8. Immunoreactivity of human mAb panel against uLAM

Other than the A194-01 IgG, the reactivity of the other mAbs described in this study against urinary LAM antigens have not been previously described. To test this, we performed a checkerboard analysis of all eight of the mAbs against ManLAM and a set of five clinical urine samples collected from HIV-1 coinfected subjects with microbiologically confirmed TB, seen at the Infectious Disease Institute at Makerere University (Suppl. Fig. 4). We included the MTX-dependent S4-20 mAb as a capture reagent, to allow comparisons with the current standard test, and in view of its enhanced binding properties, we used the A194-01 IgM as a detection reagent. Most of the antibody combinations (44/81) strongly recognized ManLAM, with S/N ratios >100. The strongest reactivity for all of the capture antibodies was provided by the S4-20 detection reagent, followed by the A194-01 IgM, while the weakest reactivity was seen for RU95-C1 and RU61-H5.

Dramatically different patterns were obtained for the clinical urine samples, for which only 6/81 of the mAb combinations gave positive signals (S/N >10) for the majority of clinical samples. The strongest signals were obtained with the A194-01 IgM detection reagent, combined with either A194-01 or RU83-A8 as capture reagents. The S4-20/A194-01 IgM pair gave very strong signals (S/N ratios of 18,476 and 900) for two of the urines, and weaker but positive signals (S/N ratios ranging from 11 to 21) for two of the three remaining urines. In contrast to its poor sensitivity against ManLAM, positive signals (S/N ratios of 61-484) were also obtained for the RU95-C1/A194-01 IgM pair for 4/5 urines, and for the RU95-C1/S4-20 combination (S/N ratios of 47-120) for 3/5 tested urine samples. The RU61-H5/A194-01 IgM pair also gave weak positive signals (S/N ratios 17-139) for 4/5 of the urines. The other combinations provided sporadic positive signals for limited numbers of urine samples.

## Discussion

The characteristics of the eight mAbs described in this study are summarized in Table 1. These antibodies were all derived from human subjects infected with *M.tb*., and the diversity of the epitopes recognized demonstrated the immunological complexity of LAM, and the range of specificities in the carbohydrate motifs of these antigens recognized by the human antibody response. The epitope specificities of the eight mAbs can be divided into four general groupings; A194-01 and RU83-A8 bound preferentially to uncapped Ara4/6-related structures, RU49-01 and RU49-02 recognized linear poly-Ara structures, P30B9, RUH5 and RU28-01 were highly specific for mannose cap-dependent structures, and RU95-C1 uniquely recognized a mannan-specific structure. The positive or negative effects of mannose capping on antibody reactivity accounted for their relative preferences for ManLAM vs. PILAM.

The cross-reactivity of S4-20 with *M.tb* LAM and LM was previously described^42^, and similar cross-reactivities between the LAM caps and LM were seen for mAbs RU61-H5, RU28-01 and RU95-C1 (but not for P30B9). The ability of these mAbs to bind to LM established that their essential epitopes consisted solely of mannose, and were not dependent on any arabinose residues. The identification of complex epitopes in LM is consistent with a report describing extended mannose branches on the mannan domain of *M. smeg* LM ^37^. RU61-H5 and RU95-C1 also recognized the mannan domain generated from LAM by endo-arabinose digestion, whereas RU28-01 and S4-20 did not (Fig. 1c), indicating differences in processing of the mannose branches between LM and LAM. The binding of RU95-C1 to the mannan domains of LAM and LM, but not with any of the mannose cap structures, is consistent with the unique recognition by this antibody of a conserved α-(1→6)-linked poly-mannose structure that is expressed in the mannan core of both LM and LAM (Fig. 2a-b).

LM is an alternate product of the PIM4 biosynthetic precursor to LAM, and consists of a phosphatidylinositol membrane anchor attached to an α-(1→6)-linked mannan core modified by various α-(1→2)-linked mannose branches, that is related to the mannan domain of LAM ^38^. LAM and LM are both essential mycobacterial envelope components that mediate pathogenicity, with accumulating evidence suggesting that LM is an equal, if not primary, driver of virulence and immune manipulation ^39,40^. The reported impact of LM on host immunity is multi-faceted: LM from from *M. chelonae* and *M. kansasii* induced TLR2/CD14-mediated TNF- and IL-8 secretion ^40^, while LM from virulent *M. tb*, but not avirulent *M. smegmatis,* potently inhibited TNF biosynthesis in human macrophages ^41^, and LMs from *M. tb* and other species induced IL-12 and TLR2-mediated apoptosis ^42,43^. In addition, the structural nuances of LM have been shown to dictate its immunological phenotype. Specifically, down-regulating the MptC mannosyltransferase disrupts α-(1→2)-mannose branching, and results in compromised cell wall integrity and altered T-cell responses ^44^, and different levels of acylation of LM were shown to mediate immune activation vs. suppression ^45^. Finally, MTX substitutions on LM enhanced its pro-apoptotic properties in macrophages while dampening pro-inflammatory cascades ^12^. Characterizing these diverse functions requires precise analytical reagents. The new information provided in this study regarding the antigenic structures of LM and the LAM mannan domain, together with well-characterized mAbs capable of differentiating these targets, provide valuable tools to fractionate distinct LM isoforms and probe the functional contributions of specific structural features.

The genetic analyses of these mAbs provides new insights into the role of antibody sequence and their function. The anti-LAM antibodies described were derived from multiple VH and VL gene families and displayed substantial sequence diversity, suggesting LAM is recognized by a genetically diverse antibody repertoire. The distribution of VH gene family usage by the antibody panel (five VH3, two VH4 and one VH1) is similar to that used by the normal healthy antibody repertoire (VH3-54.8%, VH4-23.5%, VH1-15.8%)^46^. The dominant usage of the VH3 gene family by the antibody panel is consistent with its known compatibility for carbohydrate recognition, and the other VH families have also been reported to mediate recognition of various carbohydrate antigens ^47–49^. The high level of homology and sharing of a common VH3-20/VK3-15 pairing by A194-01 and RU83-A8, despite their independent isolation from two different individuals, suggests a convergent or public antibody response to LAM, a phenomenon previously described for conserved immune responses against different microbial antigens ^50–52^. This further suggests that additional members of this family exist, which may have higher affinities for uLAM and greater efficacy for immunodetection of *M.tb* infection.

Interestingly, two mannose-reactive antibodies, P30B9 and RU28-01, contained unusually long insertions within the HCDR2 region, which make important contributions to antigen recognition. The modification of the VH4-34 CDR2 of P30B9 by a six–amino-acid insertion and an NHS→THS substitution is consistent with the reported effects of hypermutation and inactivation of a common N-linked glycosylation sequon in the VH4-34 CDR2 domain on loss of autoreactivity and development of antigen specificity ^53^. Similar insertions have previously been reported for broadly neutralizing HIV-1 antibodies that target the oligomannose-rich glycan patch of the viral envelope, where they contribute directly to mannose patch recognition on the trimeric envelope protein of the virus ^54^.

Carbohydrate-specific antibodies often possess intrinsically weak monovalent affinities for glycan epitopes, and are often expressed as higher-valent antibodies, which provides multiple contact sites for polyvalent carbohydrate-dependent targets characteristic of the surfaces of bacterial pathogens. The multi-branched structures of the arabinan domain of LAM provides for multivalent expression of epitopes dependent on both mannose-capped and uncapped poly-Ara structures, and was reflected by the strict dependence on higher-valent forms of the two Man2 cap-dependent mAbs, P30B9 and RU65-H1, for binding to LAM. This may also account for the increased sensitivity of the IgM form of A194-01 in detection of the urinary antigen.

The checkerboard analysis shown in Suppl. Fig.4 highlights the distinct immunoreactivity patterns for bacterial and clinical antigens; while most antibody combinations can efficiently detect Man-LAM, only a limited number of combinations detect urinary LAM, with A194-01 being a particularly sensitive capture and detection reagent for uLAM. Excluding the 06-013 urine, which had a particularly high level of S4-20 reactivity (S/N ratio >18,000), the strongest reactivity against the urine panel was seen for the A194-01 IgG capture and A194-01 IgM detection combination, which gave S/N ratios ranging from 1,494 to 5,284 (average signal of 2,783). The standard S4-20/A194-01 combination for the remaining four urines was considerably weaker (ranging from 9-900, average signal of 235). The second generation LAM assays currently in use all incorporate the IgG form of A194-01. The demonstration of the broader epitope specificity of the IgM form of A194-01 (Fig. 5c), and the enhanced sensitivity of this form to recognize uLAM present in multiple clinical urine samples from TB patients (Fig. 6), suggests that substituting the IgM form of A194-01 may increase the sensitivity of assays targeting clinical forms of LAM.

Also interesting was the breadth of RU95-C1 for the urine samples, despite its weak rectivity with ManLAM. The RU95-C1/A194-01 IgM combination recognized four of the five urine samples, with S/N ratio ranging from 61-484 (average = 225) for the positive samples. Although these signals were considerably weaker than those seen for the A194-01 IgG/A194-01 IgM combination, they were stronger than the corresponding signals obtained for the S4-20 capture reagent for most of these samples. In addition to high sensitivity, high specificity is also critical for a useful assay, and these experiments did not address the relative specificities of the various antibody combinations. These results suggest that further studies to screen these new combinations against larger clinical panels are merited, and may lead to new reagents and antibody combinations that improve the sensitivity of immunoassays specific for clinical forms of LAM antigens.

## Acknowledgements

This work was supported by a grant from the Bill & Milinda Gates Foundation (INV-004048) and by NIH grants R01AI152157 and R56AI171023 awarded to A.P., and by a grant from the New Jersey Health Foundation (NJHF-PC92-20) awarded to AC.

## Methods

### Study Design and Objectives

This study evaluated human monoclonal antibodies targeting *M.tb* LAM and LM, with the goal to find novel antibody combinations having higher specificity and sensitivity for immunodetection of uLAM. Objectives included characterization of antibody gene usage and level of somatic hypermutations, isotype dependence for antigen recognition, antigen specificity, epitope mapping, and performance of antibody pairs in sandwich immunoassays against clinical urine samples. Experiments were exploratory and comparative, consistent with early-stage diagnostic development studies.

### Human Participants

All methods were performed in accordance with the relevant guidelines and regulations. Plasma and PBMC were collected from subjects enrolled in the study to characterize the humoral response against TB (Pro20160000116) approved by the institutional review board of the Rutgers University. Study subjects were seen at the Waymon Lattimore Practice at Rutgers University, which provides tuberculosis patient care and TB control services in Essex and Union counties in northeast New Jersey, and all provided written informed consent prior to their enrollment. Blood was processed to obtain plasma and PBMCs within a few hours of collection, and frozen until use at −80 °C. Urine from patients diagnosed with TB and anonymized control urine samples obtained from healthy volunteers were boiled for 10 minutes, centrifuged and aliquoted and frozen at −80 °C until further use.

Additional urine samples were obtained from adults, including subjects with microbiologically confirmed tuberculosis and HIV-1 coinfections, at the Infectious Diseases Institute, Makerere University, Kampala, Uganda. Study protocols were approved by relevant institutional review boards and national ethics committees. Written informed consent was obtained from all participants before enrollment. TB-positive participants were identified with clinically confirmed diagnostic tests for latent or microbiologically confirmed active tuberculosis. Negative control urine samples were obtained from individuals without uberculosis. Urine samples were collected under standardized clinical procedures, and heat inactivated at 56°C for 1 hour and filtered with 0.2-micron filters. After centrifugation, samples were aliquoted to minimize freeze–thaw cycles, stored at −20 °C until analysis and thawed once prior to testing.

### Antigens

Purified mycobacterial lipoglycans were obtained from BEI Resources (NIAID, NIH). Antigens included lipoarabinomannan from *Mycobacterium tuberculosis* (ManLAM), *M. smegmatis* (PILAM), and *M. leprae* (LepLAM), as well as lipomannan (LM) from *M. tuberculosis*. Antigens were reconstituted according to supplier instructions and stored at −80 °C until use.

### ELISA methodologies

#### Direct ELISA for antigen binding

Binding of monoclonal antibodies to LAM and LM was assessed by enzyme-linked immunosorbent assay (ELISA). High-binding 96-well microtiter plates were coated overnight at 4 °C with purified lipoglycan antigens (2 µg/mL) diluted in carbonate–bicarbonate buffer (pH 9.6). Plates were washed with phosphate-buffered saline containing 0.05% Tween-20 (PBST) and blocked with 2% (w/v) non-fat dry milk in PBST for 1 h at room temperature. Serial dilutions of mAbs were added to the plates and incubated for 1 h at 37 °C. After washing, bound antibodies were detected using alkaline phosphatase (AP)–conjugated anti-human secondary antibodies. Plates were developed using *p*-nitrophenyl phosphate (pNPP) substrate prepared in diethanol amine buffer (1 M diethanolamine, 0.5 mM MgCl₂, pH 9.8), and absorbance was measured at 405 nm using a microplate reader.

#### Chemiluminescent ELISA for high-sensitivity detection

For enhanced sensitivity, ELISA was performed using horseradish peroxidase (HRP)–conjugated anti-human secondary antibodies. Serial dilutions of plasma or mAbs were incubated with antigen-coated plates for 1 h at room temperature. Following washing, HRP-conjugated secondary antibody was added and incubated for 1 h. Plates were developed using the Advansta ELISA Bright chemiluminescent substrate, following the manufacturer’s instructions, and signal intensity was quantified as relative luminescence units (RLU), using a 96-well luminescence plate reader.

#### Sandwich ELISA for Antigen Detection

For antigen capture assays, plates were coated overnight at 4 °C with capture antibody (10 µg/mL) in carbonate–bicarbonate buffer (pH 9.6), and washed and blocked as described above. Antigen samples were added and incubated for 1 h at room temperature, washed to remove unbound antibody, and biotinylated detector antibody (0.2 µg/mL) was added and incubated for 1 h. Following additional washes, streptavidin–horseradish peroxidase (HRP) conjugate diluted 1:2,500 in blocking buffer was added (50 µL per well), and incubated for 30 min at room temperature. Plates were washed at least of three times with PBST, developed by incubation with gentle agitation for two minutes with the chemiluminescent substrate, and signal measured as relative luminescence units (RLU) using a microplate luminometer. Each sample and condition was tested in triplicate wells, and representative averaged data from independent experiments are shown.

### Western Blot Analysis

Lipoglycan antigens were separated by SDS–PAGE and transferred onto nitrocellulose membranes. Membranes were blocked with 5% milk in PBST and incubated with primary antibodies. Bound antibodies were detected using HRP-conjugated anti-human light-chain secondary antibodies, and signals were visualized by chemiluminescence.

### Epitope mapping using synthetic glycoconjugate microarrays

MAb reactivity to LAM glycan epitopes was determined using a microarray containing 63 synthetic mycobacterial cell wall glycans, including both LAM and non-LAM structures ^46^. Microarray slides were treated with Blocker™ BLOTTO in TBS at 4 °C overnight, and incubated with mAbs at seven serial four-fold dilutions for 1 h at 37 °C, with shaking at 600rpm. After washing (10×) with 0.05% TBS-T, the slides were incubated with the secondary antibody conjugated with Cy3 at 1:1000 in BLOTTO for 40 min at room temperature. Images were scanned with a GenePix 4000 Microarray scanner system (Molecular Devices, CA), and analyzed using GenePix Pro 7.3.0.0 software to measure median pixel intensity (MPI) and neighboring background pixel intensity (BPI) of individual spots. The median fluorescence intensity (MFI), representing epitope specific mAb reactivity, was calculated using the MPI minus the BPI and averaged from the triplicate spots.

#### Enzymatic Digestion of LAM with Endo-MA1

Endo-MA1, an endo-α-1-D-arabinofuranosidase that selectively degrades the arabinan backbone while preserving the mannan core, was expressed from the pET23d expression plasmid kindly provided by Dr. Kiyotaka Fujita (Kagoshima University, Japan) and purified as previously described ^28^. Briefly, 100 ng of the plasmid was transformed into *E. coli* BL21-CodonPlus(DE3)-RIL cells (Agilent Technologies), and transformants were selected on LB agar containing ampicillin (100 μg/mL) and chloramphenicol (34/μg mL). A single colony was inoculated into 5 mL LB medium supplemented with ampicillin and chloramphenicol and cultured overnight at 37°C with shaking at 280 rpm. The overnight culture was diluted 1:100 into 150 mL auto-induction medium and the appropriate antibiotics and incubated for 20 hrs. Bacterial suspensions were lysed by sonication and lysates were clarified by centrifugation and filtered through 0.45 μm membranes. The enzyme has an attached C-terminal HisTag, and was purified by affinity chromatography using Ni²⁺-NTA resin. After washing, bound enzyme was eluted with 200 mM imidizole. Protein purity was confirmed by SDS-PAGE, and protein concentration was determined spectrophotometrically. For enzymatic digestions, ManLAM was incubated with purified Endo-MA1 for 1 hr in PBS, and enzyme was then deactivated by boiling. Digested LAM preparations were subsequently analyzed by SDS-PAGE and tested for antibody binding by ELISA and ECL assays.

#### Isolation of Anti-LAM Monoclonal Antibodies

Anti-lipoarabinomannan (LAM) monoclonal antibodies (mAbs) were isolated from human memory B cells as previously described ^9^. Peripheral blood mononuclear cells (PBMCs) were isolated from individuals with confirmed tuberculosis infection, and memory B cells were enriched using a combination of negative selection followed by positive selection with magnetic beads according to the manufacturer’s instructions. Purified memory B cells were cultured in vitro in 96-well plates at a limiting density of approximately 100 cells per well, and co-cultured with 1.5 × 10⁴ MS40L-low feeder cells per well in a total volume of 200 μL. MS40L cells were derived from murine stromal MS5 cells transduced to express human CD40 ligand (CD40L). Cultures were maintained in complete medium supplemented with a Toll-like receptor 9 (TLR9) agonist (CpG ODN 2006; 1 μM) and recombinant human cytokines, IL-2 (10 ng/mL), IL-4 (2 ng/mL), IL-10 (100 ng/mL) and IL-21 (100 ng/mL) (all from Miltenyi Biotec), to promote B cell activation, proliferation, and antibody secretion.

After 14 days of culture, supernatants were screened for LAM-specific antibodies by ELISA using purified ManLAM as antigen, and wells exhibiting positive binding signals were selected for sub-cloning. Positive cultures were serially diluted to achieve limiting dilution conditions corresponding to approximately 1–5 B cells per well, and cultured under the same conditions. After an additional 8–10 days, culture supernatants were rescreened by ELISA to confirm antigen specificity. Wells demonstrating reproducible LAM binding were further characterized to determine the LAM binding antibody isotype and light chain usage using ELISA with anti-human IgG, IgM, IgA, κ, and λ secondary reagents. Cultures estimated to contain fewer than five distinct B cell clones were selected for cDNA preperation.

#### Molecular cloning of monoclonal antibodies

Cells from selected wells were lysed, and cDNA was generated by reverse transcription PCR (RT-PCR), using the SuperScript™ IV Single Cell/Low Input cDNA PreAmp Kit (Thermo Fisher Scientific) according to the manufacturer’s protocol. Immunoglobulin genes for the heavy and light chains of the variable (V) region (Ig VH and Vκ) were amplified using random hexamer primers and then sequenced. Nested PCR was used to clone Ig genes into IgG1 heavy- and light-chain expression vectors which were co-expressed by transfection of 293 HEK cells19,20. Germline alleles were determined using the IMGT database (http://imgt.org). The V_H_ and V_L_ regions cloned from positive wells were inserted into H chain (IgG1) and L chain (κ) expression vectors and sequenced. All combinations of H and L chains were transfected into 293T cells, and H and L chain combinations reactive with LAM were selected by ELISA.

#### Phylogenetic analysis of heavy- and light-chain variable region sequences of human anti-LAM monoclonal antibodies (Suppl. Fig. 2)

Phylogenetic relationships among immunoglobulin heavy-chain variable (IGHV; panel A) and light-chain variable (IGLV/IGKV; panel B) region sequences of anti-LAM monoclonal antibodies were analyzed to evaluate sequence diversity and germline relatedness. Nucleotide sequences were aligned using Clustal Omega, and phylogenetic trees were generated using MEGA X software ^55,56^. Evolutionary history was inferred using the Neighbor-Joining method with bootstrap analysis performed using 50 replicates ^57,58^. Evolutionary distances were calculated using the Poisson correction method and are presented as the number of amino acid substitutions per site. All positions containing gaps or missing data were excluded from the analysis.

#### Checkerboard analysis of efficiency of different mAb combinations to detect urinary LAM in ECL assays

96 well luminometry plates (Greiner600) were coated with capture anitbodies at 5 ug/ml in 0.1M bicarbonate buffer for 18 h at 4°C, and then washed with PBS containing 0.05% Tween-20 and blocked with 2% non-fat dry milk (Carnation) in PBS with sodium azide for 1 h at room temperature. Plates were washed, and antigens diluted in 2% blocking buffer were added to triplicate wells of the plate. Uriines were tested at 1:2 dilutions and ManLAM at 10 ng/ml in blocking buffer. Plates were incubated for 1 hour at room temperature, and washed. Biotinylated detection antibodies diluted in 2% blocking buffer were added at 0.5 µg/ml (0.2 µg/ml for A194-01 IgM) to the appropriate wells, and incubated for 30 minutes at room temperature. Plates were washed and HRP-streptavidin (Jackson Immunoresearch), diluted to 0.4 ugm/ml in blocking buffer, was added to all the wells and incubated for 30 minutes at room temperature. Plates were washed and ELISA Bright substrate (Advansta, Inc.) was added according to the manufacturers instructions. Relative light units (RLU) were measured in a SpectraMax iD3 luminometer (Molecular Devices). To account for differences in antibody:antibody(+biotin) interactions, results were calculated as Signal : Background Ratios.

## Data availability

The antibody variable region sequences have been deposited in the GenBank database under accession numbers MC402173–MC402178. All data supporting the findings of this study are included in the article and its Supplementary Information.

## Figure legends

**Suppl. Fig. 1.** Mapping of glycan epitopes recognized by mAbs by binding assays to microarray containing synthetic Ara- and Man-containing glycans representing structural motifs present in ManLAM conjugated to a BSA carrier protein. a- Structures of synthetic glycoconjugates present in the microarray. b- Titration of binding activity of eight mAbs against the synthetic microarray.

**Suppl. Fig. 2. Phylogenetic analysis of heavy- and light-chain variable region sequences of human anti-LAM monoclonal antibodies.** Phylogenetic trees showing sequence relationships among immunoglobulin heavy-chain variable (IGHV; panel) and light-chain variable (IGLV/IGKV; panel B) regions of human anti-LAM monoclonal antibodies isolated from individuals with M.tb infection. Trees were generated using aligned nucleotide sequences of the variable regions and constructed based on sequence divergence from germline immunoglobulin genes. Branch lengths represent evolutionary distance, and numerical values at branch points indicate bootstrap support values or sequence divergence metrics. Closely clustered antibodies indicate shared sequence similarity and potential clonal relatedness, whereas more distant branches reflect greater diversification of variable region sequences.

**Suppl. Fig. 3**. Alignment of anti-LAM monoclonal antibody variable region sequences with nearest germline genes. Amino acid sequence alignments of immunoglobulin heavy-chain (upper panel) and light-chain (lower panel) variable regions of anti-LAM monoclonal antibodies with their nearest inferred germline sequences are shown. Framework regions (FR1–FR4) and comple-mentarity-determining regions (CDR1–CDR3) are indicated above the alignments according to IMGT numbering. Identical residues relative to the germline sequence are indicated by dashes, whereas amino acid substitutions are shown explicitly. Somatic mutations within framework and CDR regions demonstrate varying levels of affinity maturation among the anti-LAM antibodies.

**Suppl. Fig. 4.** Checkerboard analysis of 8 human LAM-specific mAbs and S4-20 for ManLAM and 5 patient urine samples. Results are presented as single:noise ratios, with values >10 considered to be positive signals.

**Suppl.Table 1. Immunoglobulin heavy- and light-chain variable rgion gene usage and somatic hypermutation profiles of human anti-LAM monoclonal antibodies.** Summarized are the immunoglobulin variable gene usage, framework region (FR) and complementarity-determining region (CDR) nucleotide lengths, and somatic hypermutation (%SHM) frequencies of the heavy-chain (top panel) and light-chain (bottom panel) variable regions of eight human anti-LAM monoclonal antibodies. Heavy-chain variable (IGHV), diversity (IGHD), and joining (IGHJ) gene segments, as well as light-chain variable (IGKV/IGLV) and joining (IGKJ/IGLJ) gene usage, were assigned by sequence alignment to germline immunoglobulin genes. Nucleotide lengths and percent somatic hypermutation are shown for framework regions (FR1–FR3) and complementarity-determining regions (CDR1–CDR3). Overall variable region somatic hypermutation frequencies for heavy (IGHV) and light (IGLV) chains are indicated in blue. Red text highlights CDR-specific features.

