## Supplementary figures and images for "Novel human monoclonal antibodies with enhanced sensitivity for lipoarabinomannan antigens present in urines of TB patients"

### Supplemental Figures

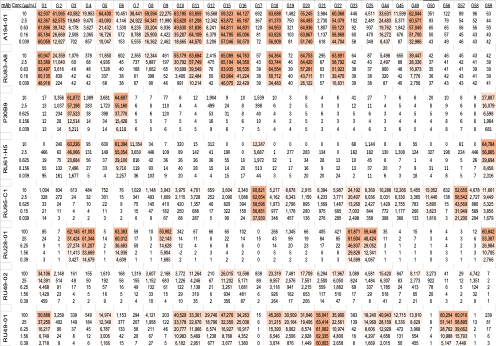
Suppl. Fig 1

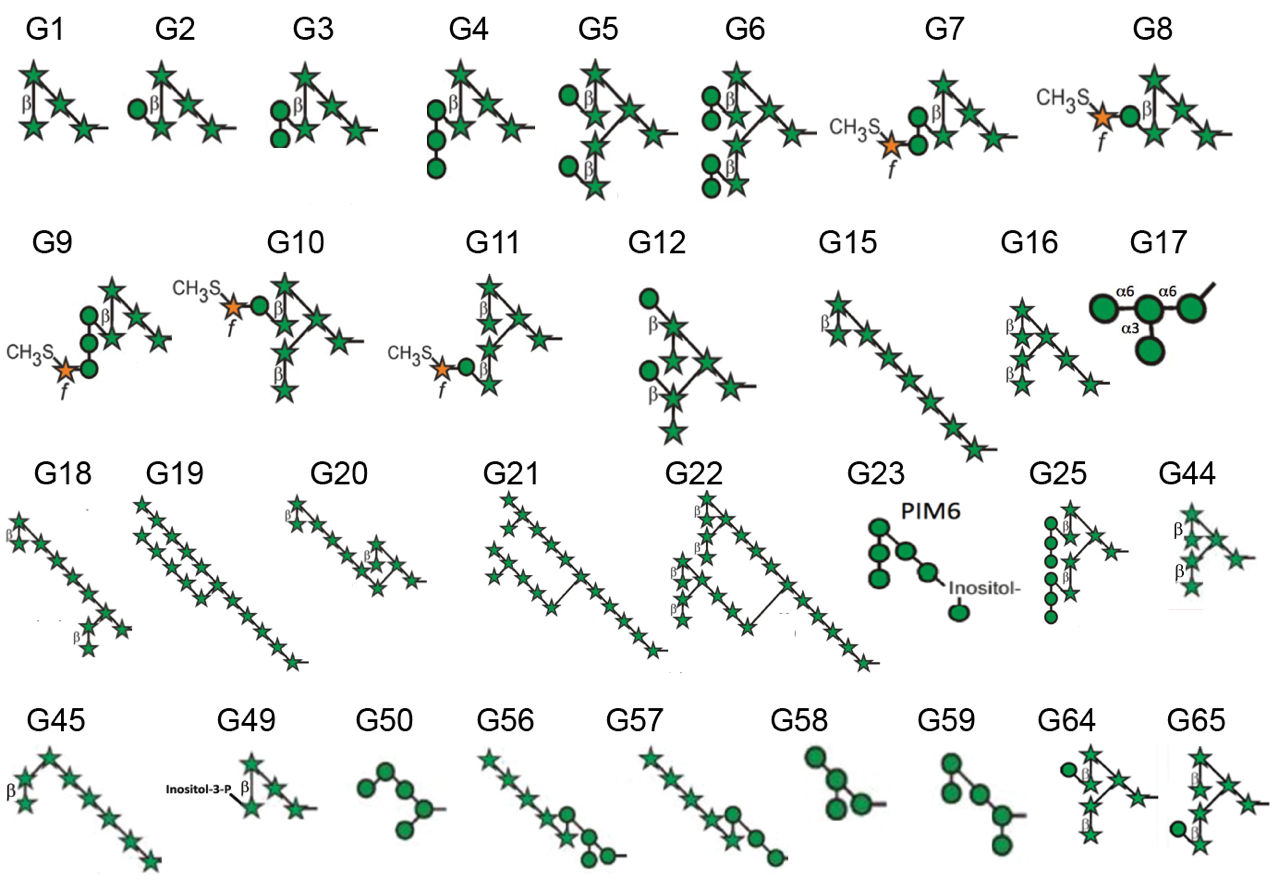
a

b

**Suppl. Fig. 2**

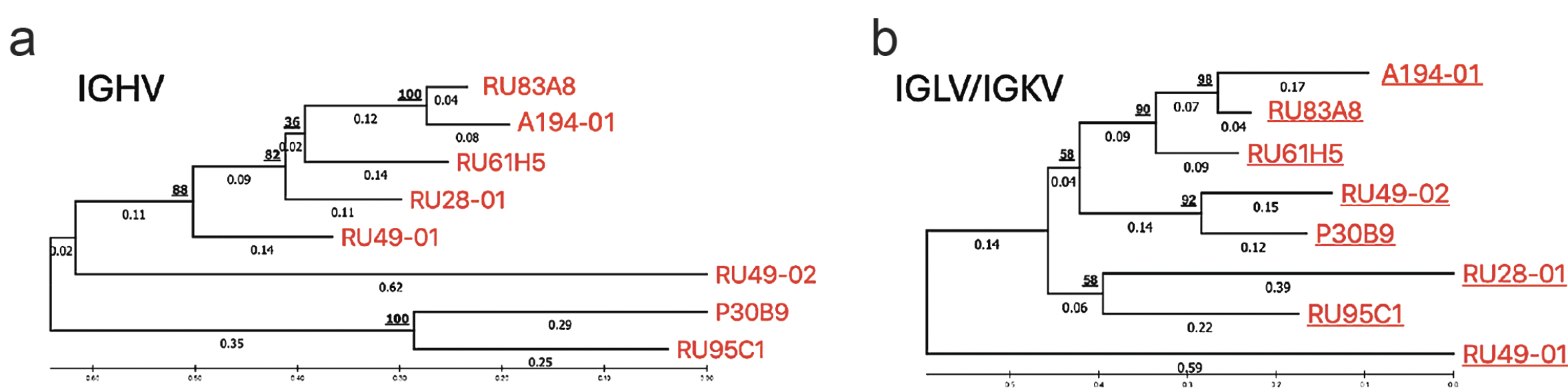

.

**Suppl. Fig. 3**

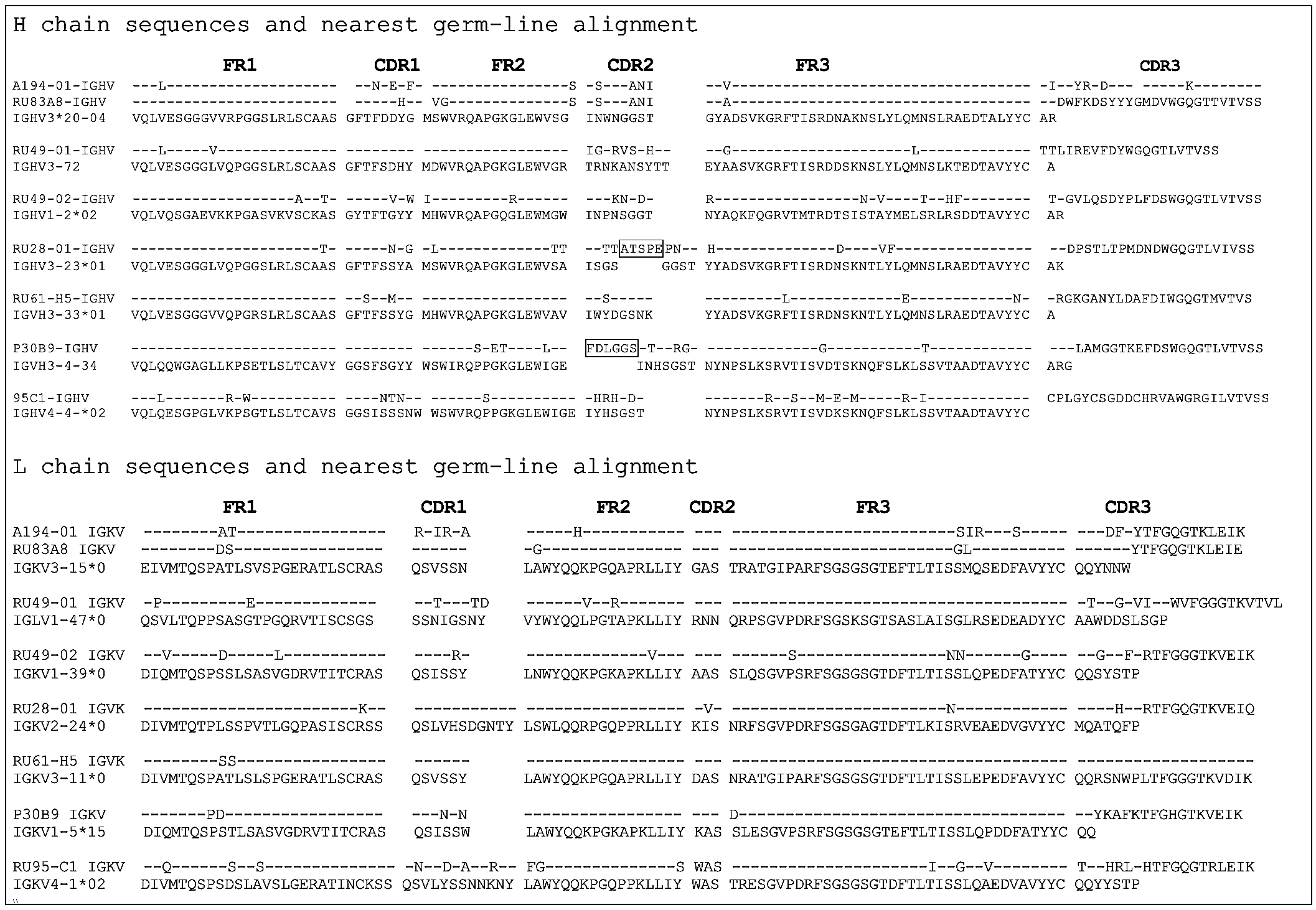

**Suppl. Fig. 4**

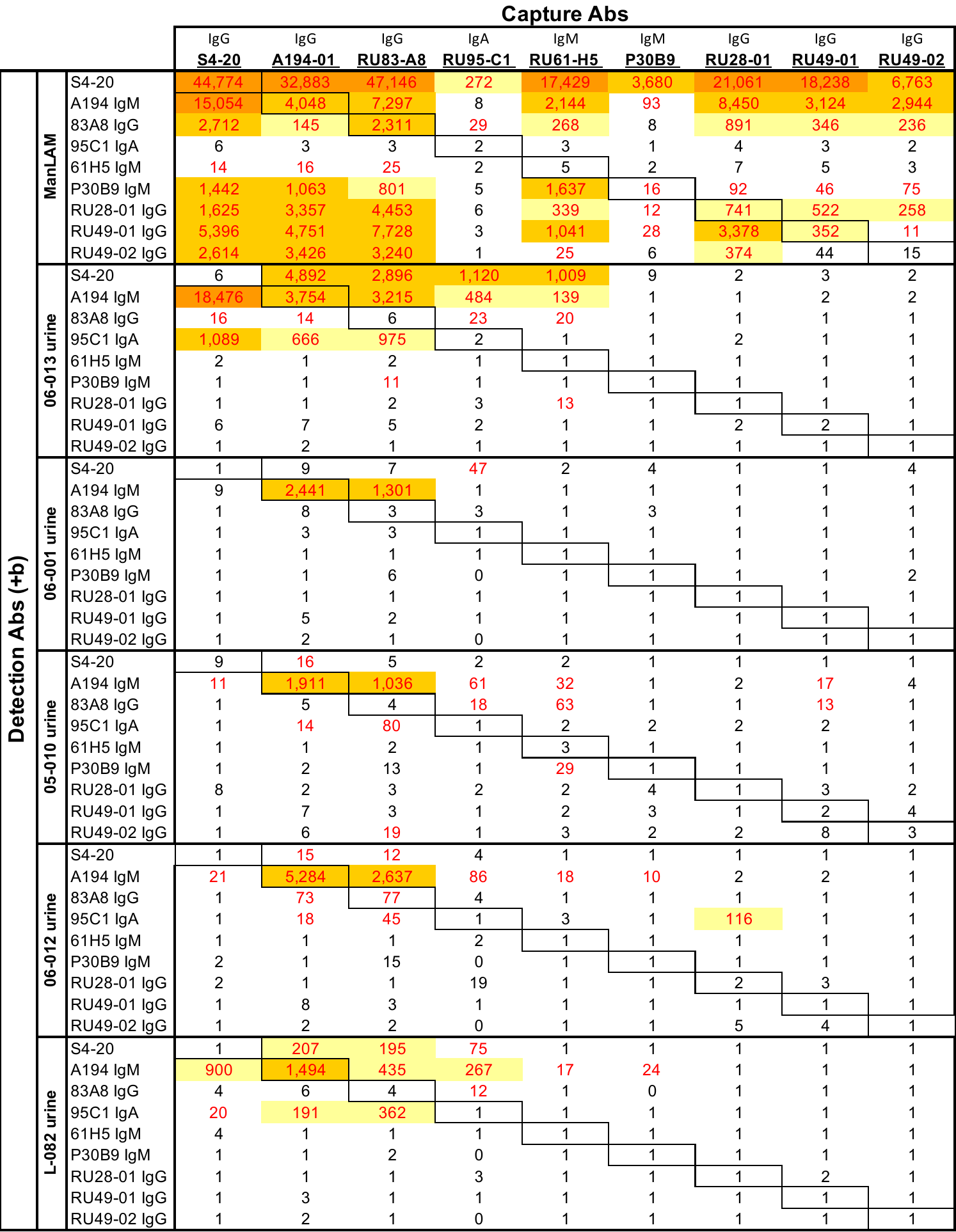

### Supplemental Tables

**Suppl. Table 1.**

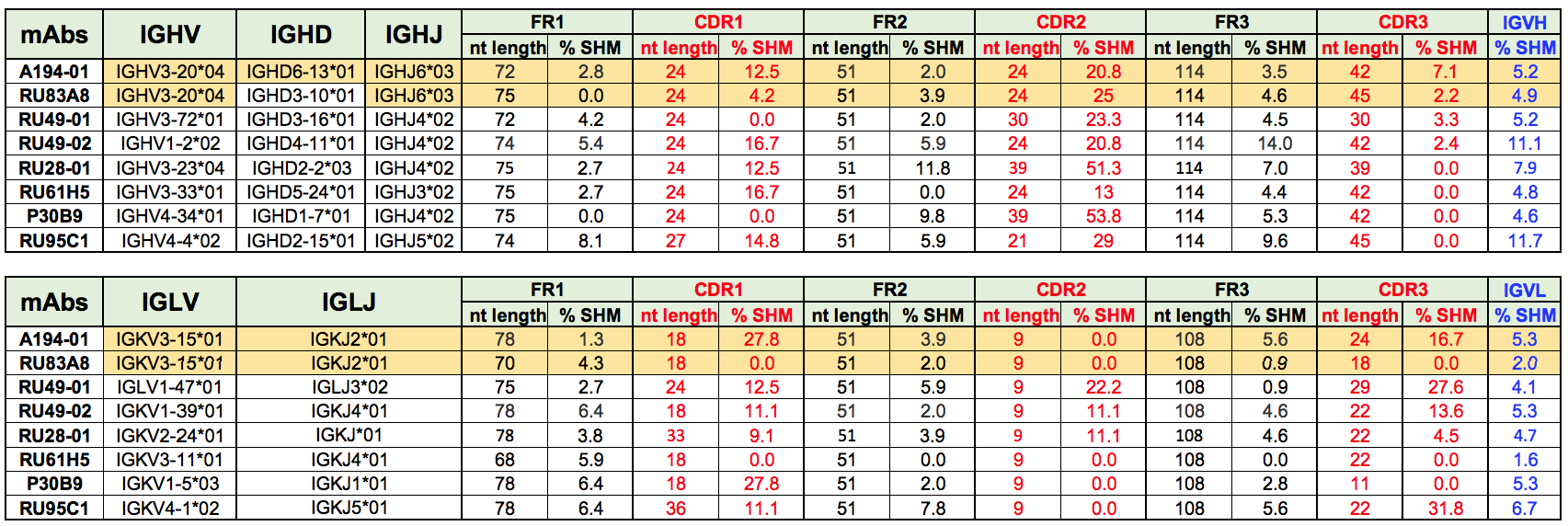
